# An integrated Asian human SNV and indel benchmark combining multiple sequencing methods

**DOI:** 10.1101/759043

**Authors:** Chuanfeng Huang, Libin Shao, Shoufang Qu, Junhua Rao, Tao Cheng, Zhisheng Cao, Sanyang Liu, Jie Hu, Xinming Liang, Ling Shang, Yangyi Chen, Zhikun Liang, Jiezhong Zhang, Peipei Chen, Donghong Luo, Anna Zhu, Ting Yu, Wenxin Zhang, Guangyi Fan, Fang Chen, Jie Huang

## Abstract

Precision medicine of human requires an accurate and complete reference variant benchmark for different populations. A human standard cell line of NA12878 provides a good reference for part of the human populations, but it is still lack of a fine reference standard sample and variant benchmark for the Asians. Here, we constructed a stabilized cell line of a Chinese Han volunteer. We received about 4.16T clean data of the sample using eight sequencing strategies in different laboratories, including two BGI regular NGS platforms, three Illumina regular NGS platforms, two linked-read libraries, and PacBio CCS model. The sequencing depth and reference coverage of eight sequencing strategies have reached the saturation. We detected small variants of SNPs and Indels using the eight data sets and obtained eight variant sets by performing a series of strictly quality control. Finally, we got 3.35M SNPs and 349K indels supported by all of sequencing data, which could be considered as a high confidence standard small variant sets for the studies. Besides, we also detected 5,913 high quality SNPs located in the high homologous regions supported by both linked-reads and CCS data benefited by their long-range information, while these regions are recalcitrant to regular NGS data due to the limited mappability and read length. We compared the later SNPs against the public databases and 969 sites of them were novel SNPs, indicating these SNPs provide a vital complement for the variant database. Moreover, we also phased more than 99% heterozygous SNPs also supported by linked-reads and CCS data. This work provided an integrated Asians SNV and indel benchmark for the further basic studies and precision medicine.

## Introduction

Thousands of human genomes are now available and whole genome sequencing (WGS) is likely to become a routine part of medical care in many countries. WGS data allows the identification of genetic changes associated with disease and paves the way for precision medicine, medical care customized according to the genetic make-up of a patient [1]. Diseases are often associated with particular single nucleotide variants (SNVs), or insertion or deletion events (indels) [2, 3]. In order to fully capitalize on the vast genome data generated, reference genomes are required to allow genome comparisons and benchmarking of new sequencing technologies and analysis methods. The current human reference genome (NA12878) is from a Caucasian from the U.S. state of Utah. Significant insights have been gained from NA12878, but it is appreciated that reference genomes from additional populations are needed [4]. Several Asian genomes are now available from individuals of Chinese [5], Korean [6] and Pakistani [7] descent. However, the majority of these studied used next-generation sequencing (NGS) platforms to generate short reads and could not resolve SNVs and indels located in complex regions. For example, targeted DNA-HiSeq [8] identified 1,281 SNVs in 193 genes in the Asian reference sample YH that could not be detected in the original study [5].The 193 genes are associated with hereditary diseases with a higher incidence in the Chinese population, a clear example of the need for high quality reference genomes in addition to NA12878 [7]. It is now apparent that a combination of long read, short read, and linked-read sequencing is required to fully characterize human reference genomes[9]. Herein, we generated an Asian SNV and indel benchmark genome by combining diverse short and long read sequencing platforms, an approach which could balance the systematic sequencing bias of different platforms.

## Results

### Sequencing and quality control

To develop a represented Asian high-quality genotype call sets, we recruited a Han Chinese volunteer from Beijing City (Research ethics ID: XHEC-C-2019-086, HJ). We sequenced this individual using five frequently-used NGS short-read sequencing platforms (BGISEQ-500, MGISEQ-2000, NextSeq-CN500, NextSeq550Dx and NovaSeq6000; three technical replicates), single tube long fragment read (stLFR) sequencing[10], 10X Genomics Chromium linked-read sequencing[11], PacBio single molecule real-time circular consensus sequencing (SMRT CCS) long-read sequencing[12] and Oxford Nanopore MinION sequencing [13]. After processing (**Figure 1**), we generated 3.12Tb high quality general NGS data for HJ totally. This included an average coverage of 86.58× from 2×100bp reads on two BGISEQ-500 sequencers and 60.07× from 150bp reads on three Illumina sequencers. We obtained 250.78 Gb (∼51.97×) stLFR data with a molecular length of 117,499 bp, 277.60 Gb (∼84.7×) 10X Genomics Chromium data with a molecular length of 191,294bp, and 77.23 Gb (∼24.4×) PacBio CCS data with mean read length of 12.09 kb. For the general NGS data, 99.88% of raw reads could be mapped to the human reference genome (hs37d5) with coverage of 99.92% and 85.75% of mapped reads were uniquely mapped reads. For the stLFR data and 10X Genomics Chromium data, we aligned 99.35% and 99.71% of them against the reference genome with 98.86% and 98.90% coverage, respectively. For the CCS reads, it could be unambiguously mapped to reference genome at 93.18% coverage (**Figure S1, Table S1**).

**Figure 1.**
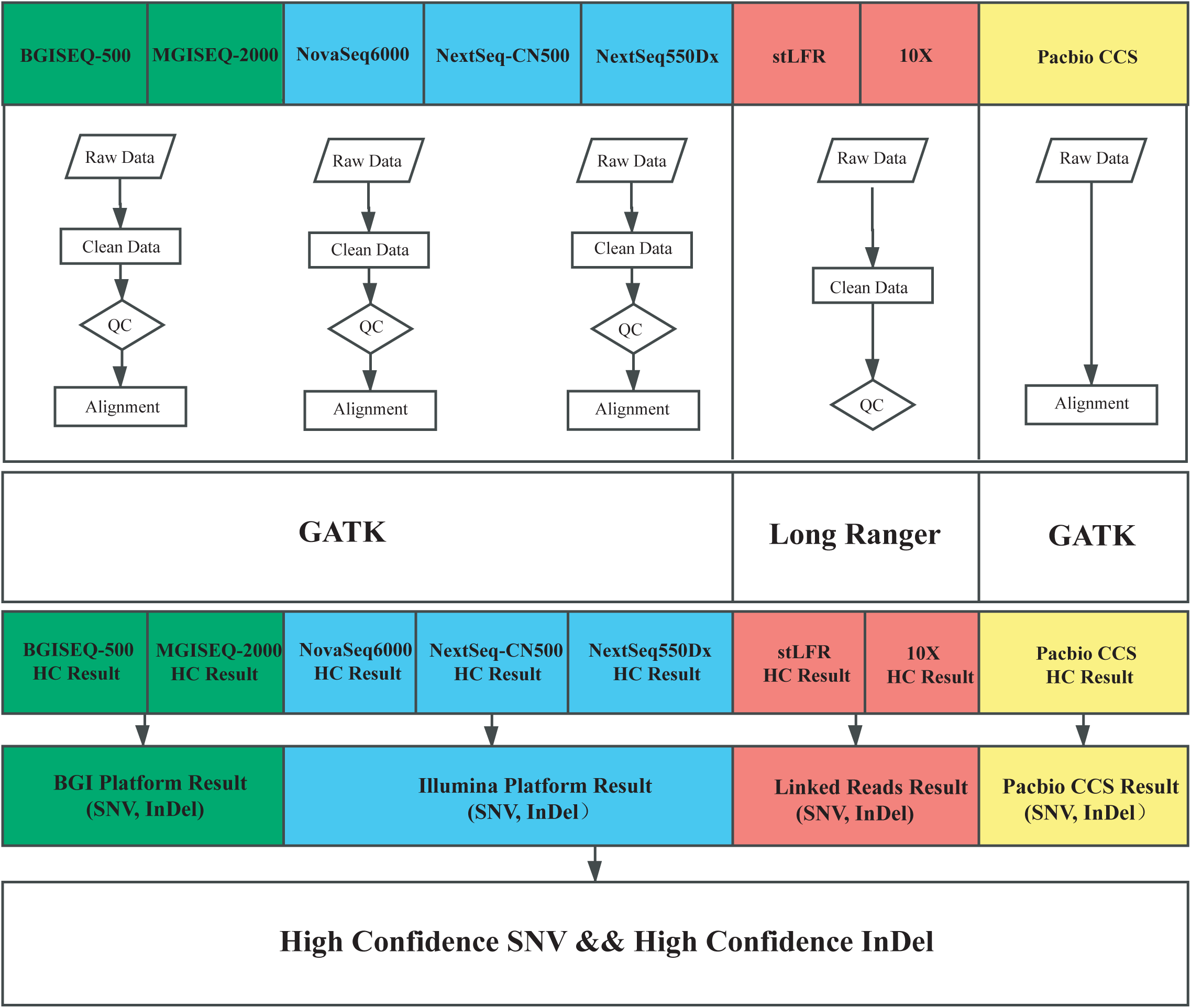
Overview of variation calling pipeline. The major steps included data filtering, alignment, variation calling, and integrated analysis.

### SNV and indel detection

To find the saturated sequencing depth of the different platforms, we hierarchically detected SNVs and indels by randomly extracting alignment results from the bam by picard. We found that 30× depth sequencing ensured a consistent rate of uniquely mapped reads (∼99%) and number of SNVs (∼3.84 M) and indels (∼897 K) (**Figure 2**, **Figure S2**).

**Figure 2.**
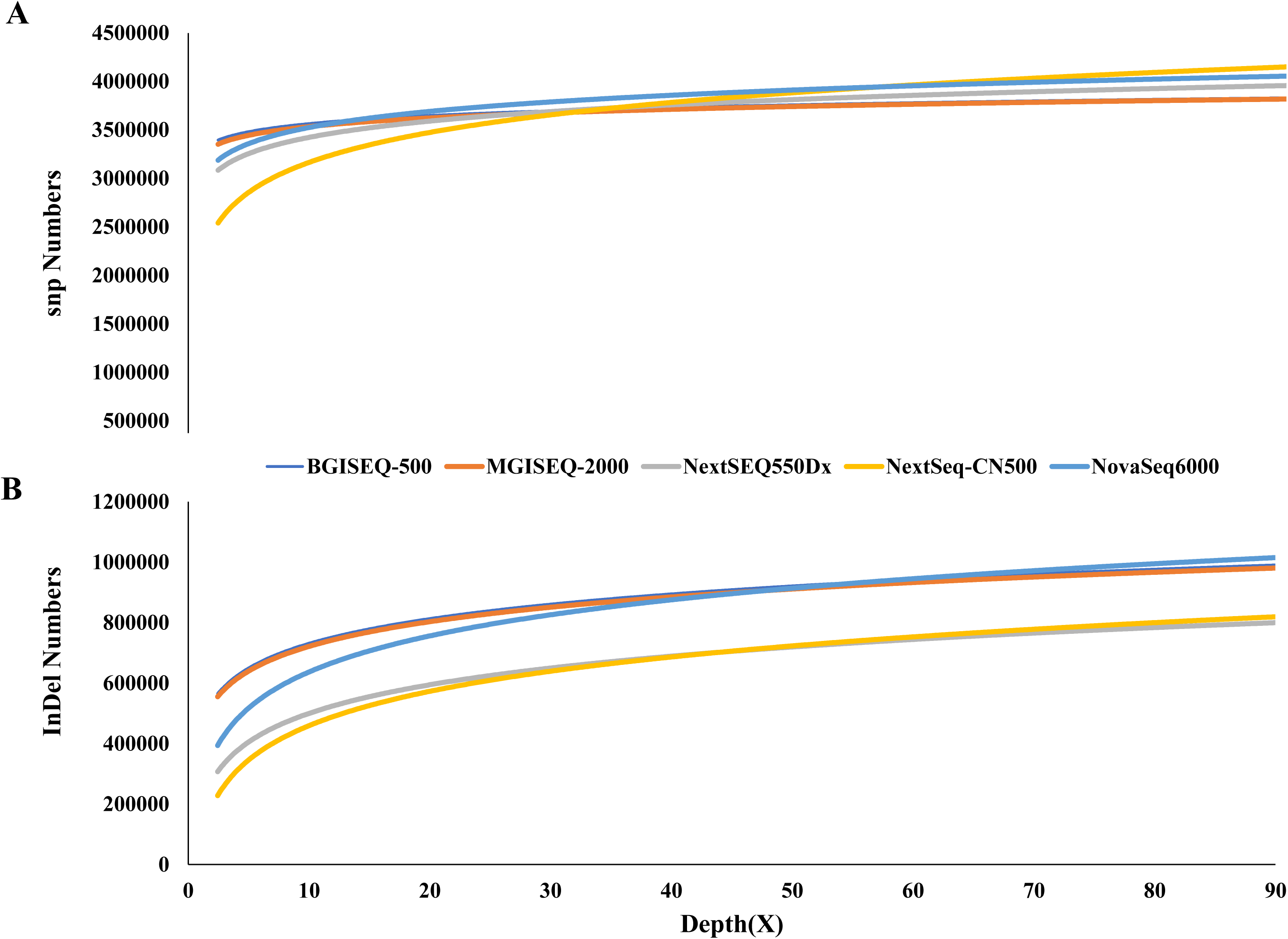
Saturation analysis. The relationship between SNPs(A)/indels(B) and depth, with the X axis for sequencing depth and the Y axis for the number of SNPs/indels detected.

We also evaluated the consistency of BGI and Illumina short sequence reads generated from short-insert libraries on the same and different instruments. We found 3.26%, 95.49%, and 1.25% of SNVs could be detected, suggesting that the choice of short-read sequencing platform introduces little bias (**Figure S3**). Nevertheless, despite an adequate sequencing depth, ∼33.62 Mb of the genome could not be resolved by short-read BGISEQ-500 and Illumina data (**Figure 3**, **Table S2**). These regions were assigned into 51,612 blocks, with an N50 of 3,942 bp, and correspond to highly homologous regions (HHRs), of which were previously reported as recalcitrant to short-read NGS sequencing[14]. Interestingly, 73.3%, 65.41% and 68.53% of these HHRs were accessible by stLFR, 10X Genomics Chromium, and PacBio SMRT CCS data, technologies which benefits from barcoding information of linked-reads or long reads (**Table S3**). We detected 3.87M, 3.47 M, and 3.80M SNPs, along with 822K, 721K, and 797K indels using stLFR, 10X Genomics Chromium, and PacBio SMRT CCS data, respectively (**Table S1**). We next wished to characterize SNVs and indels in the HJ sequencing data that could not be mapped to the Caucasian reference genome, even with long-read sequencing data. We focused on HHRs and uniquely mapped regions (UMRs).

**Figure 3.**
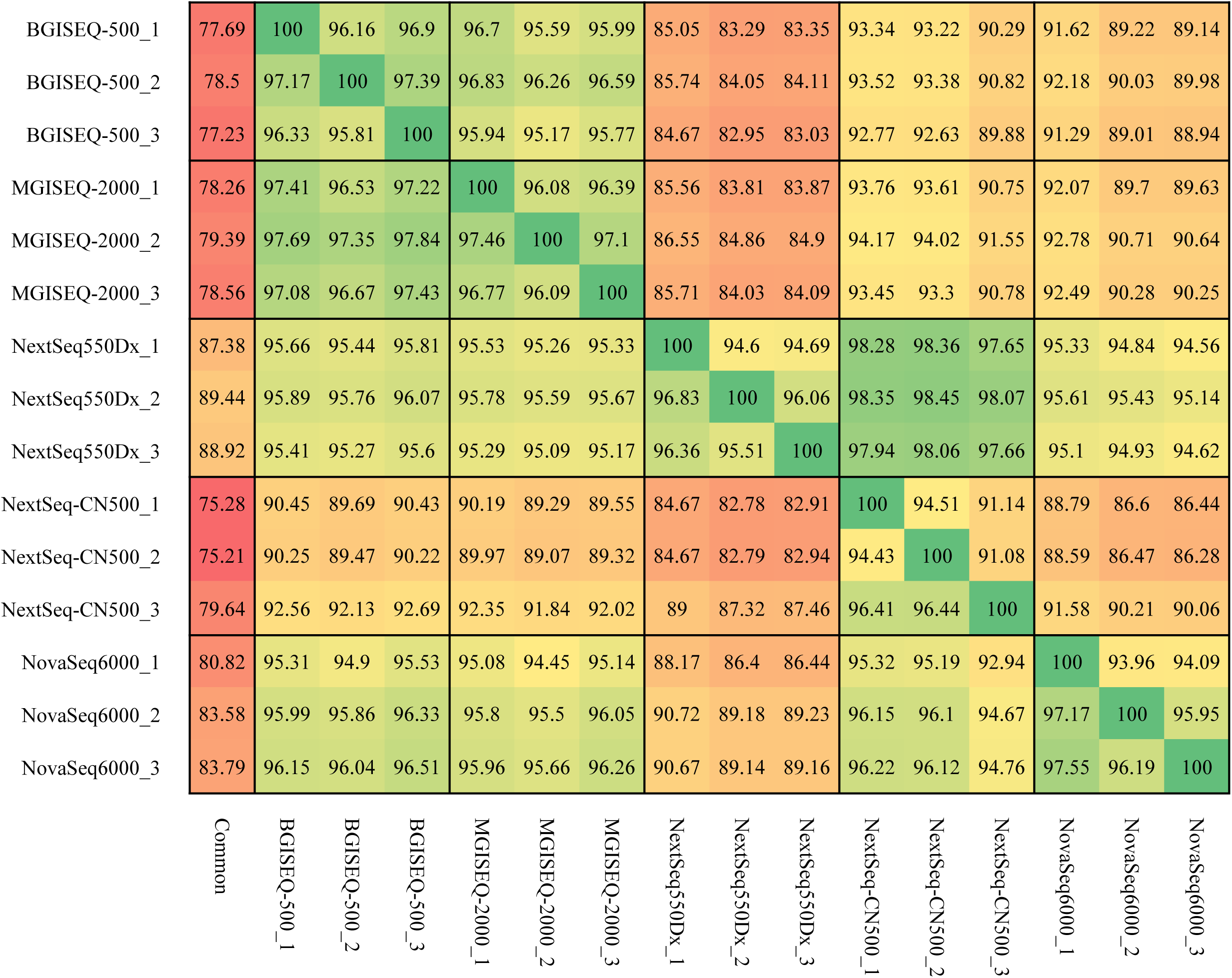
Uncovered region by NGS in each sequencing platform.

### Consistence of SNPs and indels in UMR

In the uniquely mapped regions (UMRs) 1,712,393 SNPs and 186,641 indels could be detected by all eight sequencing methods. This is less than the average number of variations (∼3.72M SNPs and ∼859 K indels). Unexpectedly, 10X Genomics Chromium missed ∼1.63M SNPs which could be detected by all of the other methods (**Figure S4**). We, therefore, excluded this data in the further analysis, retaining ∼3.35M high quality common SNPs supported by seven sequencing methods. PacBio SMRT CCS detected 234.46K specific SNPs and 240.74K specific indels; stLFR 210.45K and 223.25K; DNBSEQ-NGS 11.78K and 71K; and Illumina-NGS 5.57K and 1.98K (**Figure 4**, **Figure 5**,). We compared the SNP quality distribution between specific SNPs and whole SNPs and found that the quality of the majority of specific SNPs were lower than whole SNPs, likely stemming from sequencing method bias Interestingly, PacBio SMRT CCS and stLFR consistently resulted in high quality variant calls (**Figure S5**).

**Figure 4.**
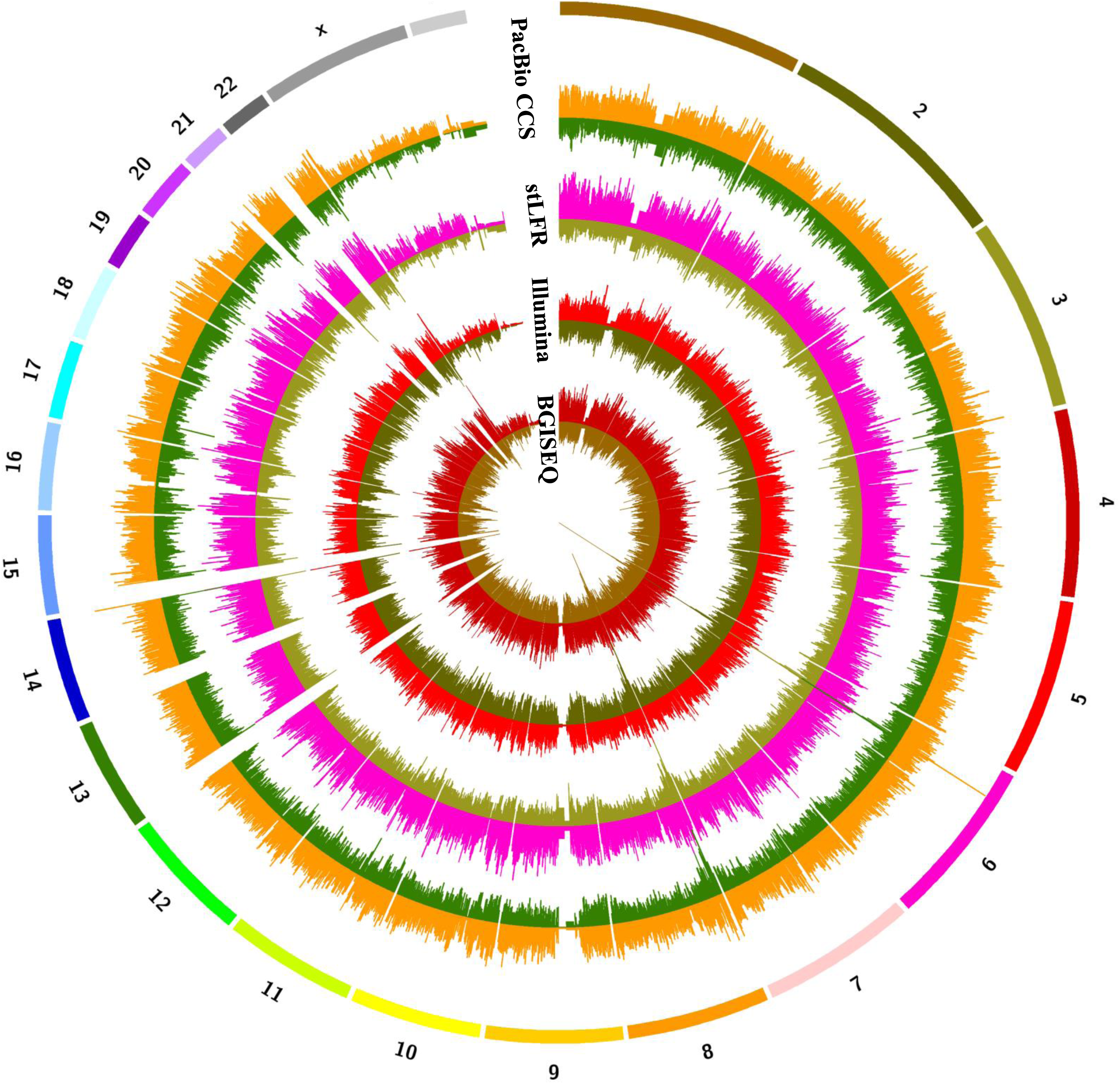
Density maps of sequencing platforms SNP and indel variations. From inside to outside circles are DNBSEQ-NGS, Illumina-NGS, stLFR and Pacbio CCS respectively, Window =1000000bp, Inside and outside are SNP and indel.

**Figure 5.**
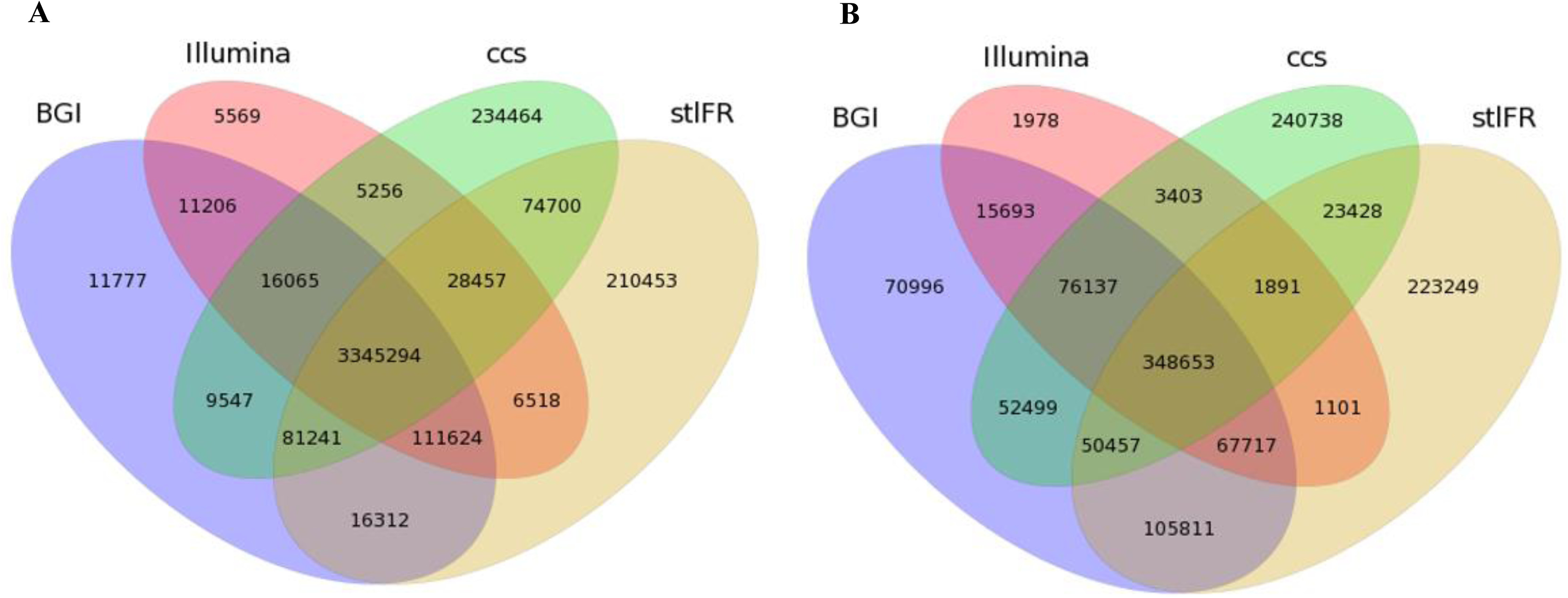
Consistency analysis: BGI regular NGS platforms, Illumina regular NGS platforms, two linked-read libraries, and PacBio CCS mode SNP(A) and indel(B) consistency analysis.

### Accessibility of SNPs and indels in HHRs

A total of 74.7K SNPs and 23.4K indels could be called by both stLFR and PacBio SMRT CCS data but not by the five short-insert library, short-read methods. These variants, nonsynonymous, were located on 129 genes and were significantly enriched for the gene ontology (GO) categories olfactory receptor activity, IgG binding, transmembrane signaling receptor activity, G protein-coupled receptor activity, molecular transducer, and signaling receptor activity pathways. We speculate that these variants are associated with immune disease in Chinese population. Among all special SNPs, 7.9% (5,913/74,717) located in HHRs, with 69 SNPs in coding regions and 19 SNPs in UTR regions. We also performed function enrichment analysis, revealing three genes significantly enriched in blood antigen-related or immune response (LILRB3, RHD, and RHCE) pathways involved into immune response diseases.

Highly homologous or repetitive regions on the genome, NGS is difficult to fully cover due to its read length, which may lead to false negative of mutations, but stLFR and CCS perform well. Complex genes are hard to be covered by NGS platforms, while linked-reads method and long reads sequences platforms do well in detecting the regions. For example, IGV shows a typical gene NBPF4, who is a member of the neuroblastoma breakpoint gene family (NBPF) which consists of dozens of recently duplicated genes primarily located in segmental duplications on human chromosome 1 (**Figure 6**).Another gene is NAIP which is part of a 500kb reverse replication on chromosome 5q13, contains at least four repeated elements and genes, and making it easy to rearrange and delete. The repeatability and complexity of the sequences also make it difficult to determine the organization of this genomic region. It is thought that this gene, modifier of spinal muscular atrophy, is a mutation in a neighboring gene SMN1. Variations detected on NAIP for NGS platform are relative small and nearly included in linked reads and long reads platforms (**Figure S6**). In addition to the genes mentioned above, there is XAGE2 (**Figure S7**), and other genes.

**Figure 6.**
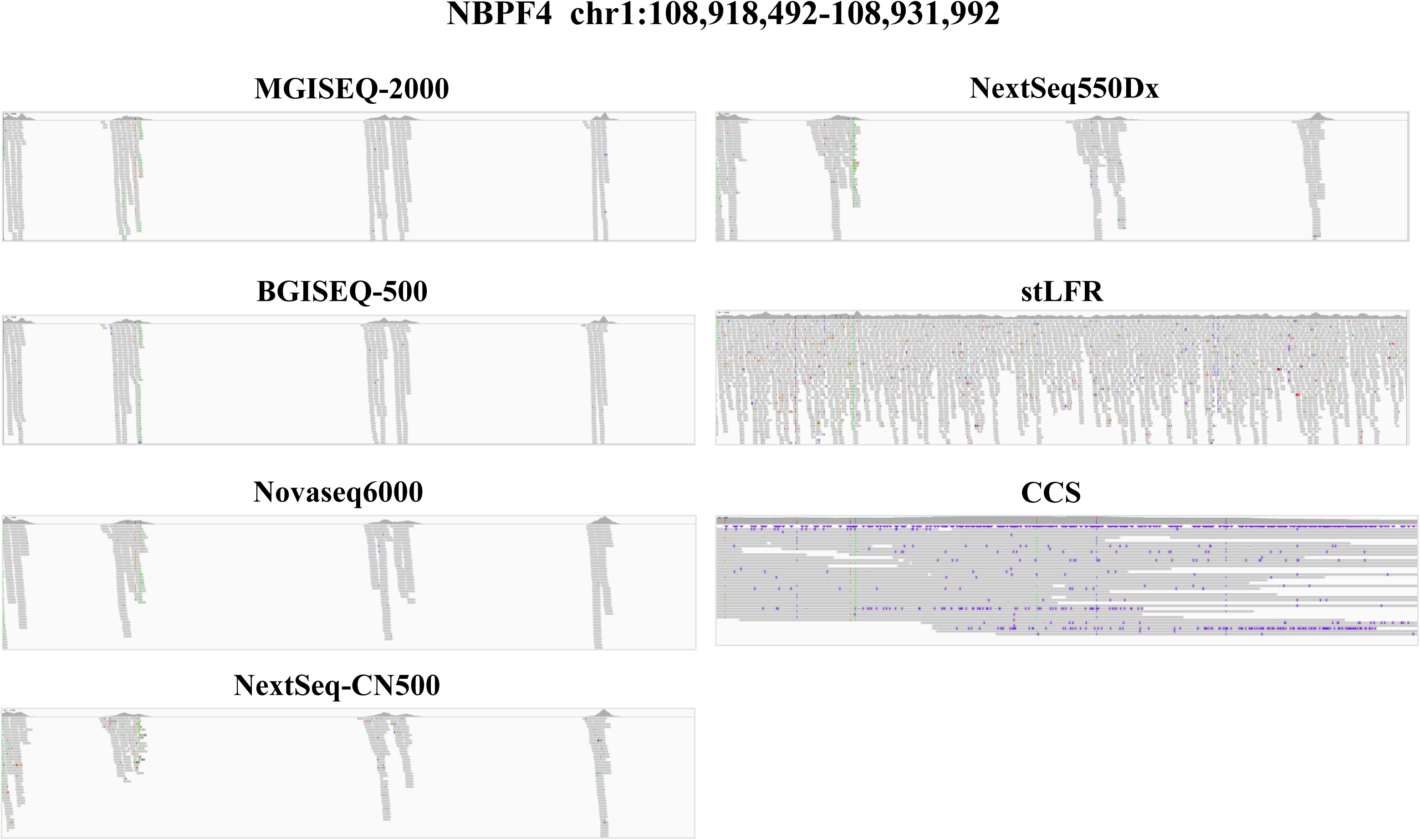
Depth and coverage of NBPF4 gene in HHRs.

### Haplotype phasing small variants

Human genomes are diploid, with chromosome pairs from each parent. However, most paired-end reads cannot assign variants to a particular chromosome, resulting in a combined haplotype (genotype) [15]. The popular NGS sequencing technology is all about shuffling sequences together for sequencing. After sequenced, we cannot directly distinguish which of these sequences is the parent source. It is only after phasing that we are able to make this distinction. Phasing is strongly correlated with functional interpretation of genetic variation. Therefore, due to the BGI and Illumina short sequence reads generated from short-insert libraries, we using long-range information from PacBio SMRT CCS and stLFR data to phasing, 99.63% and 99.91% of heterozygous SNPs could be phased into 19,584 and 1,262 blocks, respectively. Of these, 1.96 M were shared, with a phasing N50 of more than 11.26 Mb and 388.5k. What’s more, some of chromosomes (such as Chr5 and Chr6) were almost completely phased (**Table 1**). According to the results of phasing, stLFR data performed better, so we can be assumed that the long range reads may a good choice in phasing process.

## Discussion

Gene sequencing is an important part of precision medical, widely used in detection and diagnosis of various diseases, and brought potential benefits to patients. However, the NGS also some deficiencies, such as short reads, structure mutation detection, especially about the detection of HHR area will miss part of the results, thus caused by false negatives. There is currently a lack of a standard data set that represents Asian populations due to ethnic differences. In this paper, a Han Chinese adult male was recruited and 8 sequencing platforms were used to detect and compare SNV and indel. Finally, we identified a standard data set which contains 3.35M SNPs and 349K indels.

We compared and contrasted eight sequencing platforms to generate an Asian human SNV and indel benchmark. Unexpectedly, and found that 10X Genomics Chromium data did not correlate well with data generated by the other platforms, the reasons for this are not clear. However, a total of 3.35M high quality SNPs were supported by seven other methods, while linked-read stLFR and long-read PacBio SMRT CCS resolved an additional 74.7K SNPs in highly homologous regions, providing a comprehensive small variation benchmark of an Asian human. stLFR and CCS can be well supplemented and improved on the basis of NGS results.

In summary, NGS results will miss some mutations in the HHR region. By adding analysis results of stLFR and CCS platforms, standard data sets and high confidence regions that are considered relatively reliable can be obtained. This data set can be well used for further study. In order to improve the data set, it may be necessary to add samples and analysis methods for integrated analysis.

## Methods

### Sample collection

This study was carried out in accordance with relevant guidelines and regulations, in line with the principles of the Helsinki declaration[16] and was approved by the institutional review committee (IRB) of BGI. In this experiment, cell line genomic DNA was prepared from the National Institutes for food and drug Control (NIFDC), and it contained 10μg per tube. Used Qubit 3.0 to quantified the genomic DNA and agarose gel to make sure the genomic DNA molecular was not substantially degraded.

### Library and sequencing

NGS library construction adopts the normal NGS construction process. The difference between BGISEQ-500 and Illumina platform is that the former involves rolling amplification while the latter use PCR amplification technology. In particular, the DNBSEQ library protocol contain three steps: including making DNA nanoballs (DNBs), loading DNBs, and sequencing. Single tube long fragment read (stLFR) library construction physically breaks the DNA into fragments of about 50Kbps, and then use Tn5 transposase for library construction, so that each identical fragment bears the same barcode[10], while 10X Genomics Chromium library construction uses microdroplets where, after the ligation step, PCR is performed and the library is ready to enter any standard next generation sequencing (NGS) workflow.

Large-insert single molecule real-time circular consensus sequencing (SMRT CCS) library preparation was conducted following the Pacific Biosciences recommended protocols[17]. In brief, a total of 60μg genomic DNA was sheared to ∼20kb targeted size by using Covaris g-TUBEs (Covaris). Each shearing processed 10μg input DNA and a total of 6 shearings were performed. The sheared genomic DNA was examined by Agilent 2100 Bioanalyzer DNA12000 Chip (Agilent Technologies) for size distribution and underwent DNA damage repair/end repair, blunt-end adaptor ligation followed by exonuclease digestion.

### NGS data preprocess

Data filter: SOAPnuke (version 1.5.6) was used to pre-process the 15 NGS data by removing reads with (1) adaptor contaminations, (2) more than 10% low-quality bases (quality < 10), (3) more than 10% N bases.

Mapping and variant calling: All NGS reads were mapping to the human reference genome (hs37d5) using BWA 0.71.5 [18] (an in-house Apache Hadoop version). The Genome-Analysis-ToolKit (GATK) 2.3.9-lite [19] (an in-house Apache Hadoop version) was used for variant calling from BAM files with HaplotypeCaller v2.3.9-lite.

### Saturation of NGS data

Picard (version 2.18.9) was used to down-sample BAM files from 10× to the maximum depth in a 10×-step for each NGS data. Next, MegaBOLT (version 1.15) was used for variant calling from down-sampled BAM files. SNPs were hard-filtered using “QD < 2.0 || FS > 60.0 || MQ < 40.0 || MQRankSum < −12.5 || ReadPosRankSum < −8.0” and Indels were hard-filtered using “QD < 2.0 || FS > 200.0 || ReadPosRankSum < −20.0”.

### Uncovered region of NGS data

For each NGS data, any block with approximate read depth (DP) ≥ 5 were extracted from gVCF as a covered region. The uncovered regions of each NGS data were built by subtracting the covered regions from the human genome by BEDtools (v2.16.2). Meanwhile, the common uncovered regions of NGS data were built by subtracting the union of covered regions in all 15 NGS data from the human genome.

### Linked reads Mapping

The output files (FASTQ) of the linked-read sequencing methods stLFR and 10X Genomics Chromium are similar, enabling the use of the 10X Genomics Long Ranger software after converting stLFR barcodes to a Chromium compatible format. We used SOAPnuke 1.5.6 to filter out low quality and adapter reads. Clean reads were mapped and phased using the Long Ranger 2.1.2 wgs model. Briefly, de-multiplexed FASTQ files from were de-duplicated and filtered and phased SNPs, indels were called. SNP and indel information were parsed from the final VCF file using GATK SelectVariants.

### Pacbio data process

PacBio single molecule real-time circular consensus sequencing (SMRT CCS) have low base error rates, providing both highly-accurate variant calls and long-range information needed to generate haplotypes. We used the pbmm2 (version 1.0.0) alignment tool to map reads to the hs37d5 human reference genome, with the parameter --preset CCS --sample HJ –sort. GATK HaplotypeCaller was used to call SNVs and small indels. Different values of the HaplotypeCaller parameter --pcr-indel-model and VariantFiltration parameter --filter-expression were considered, setting the minimum mapping quality to 60 and using allele-specific annotations (--annotationgroup AS_StandardAnnotation) and --pcr_indel_model AGGRESSIVE. SNVs and short indels were filtered using GATK VariantFiltration with -- filter_expression of AS_QD < 2.0. Longer read lengths improve the ability to phase variants, as tools like WhatsHap demonstrate for PacBio reads [17].

## Data resource access

The sequence data from this article can be found in the CNSA databases under the following accession numbers: CNP0000091.

## Supporting information

None

## Acknowledgments

This work was supported by Grants from National Key Research and Development Program of China (2017YFC0906501, 2019B020226001). We thank Dr. Xin Liu for his guidance on the project and technical assistance.

## Competing Interests

Competing interest statement: The author denies that he has any intention to obtain any financial interests.

**Table 1**. **Haplotype phasing small variants.**

