## Supplementary material for "An integrated Asian human SNV and indel benchmark combining multiple sequencing methods": None

**Supplementary Figure S1. Sequencing depth, duplicate and mapping coverage.**

**Supplementary Figure S2. Saturate analysis for SNP, InDel and coverage for NGS platforms.**

**Supplementary Figure S3**. **Consistency of SNPs from BGI and Illumina short sequence reads.**

**Supplementary Figure S4. Consistence of SNPs and indels in UMR.**

**Supplementary Figure S5. All and platform unique SNPs quality distribution for long fragment and sort fragment platforms.**

### Supplementary Figure S6. IGV views NAIP gene for each platforms.

### Supplementary Figure S7. IGV views XAGE2 gene for each platforms.

**Supplementary Table S1-1. Data information of each platforms.**

**Supplementary Table S1-2. Data information of each platforms.**

**Supplementary Table S1-3. Data information of each platforms.**

**Supplementary Table S2. Statistics of NGS uncovered regions.**

**Supplementary Table S3. Phasing statistic for PacBio SMRT CCS, stLFR and 10X Genomics Chromium.**

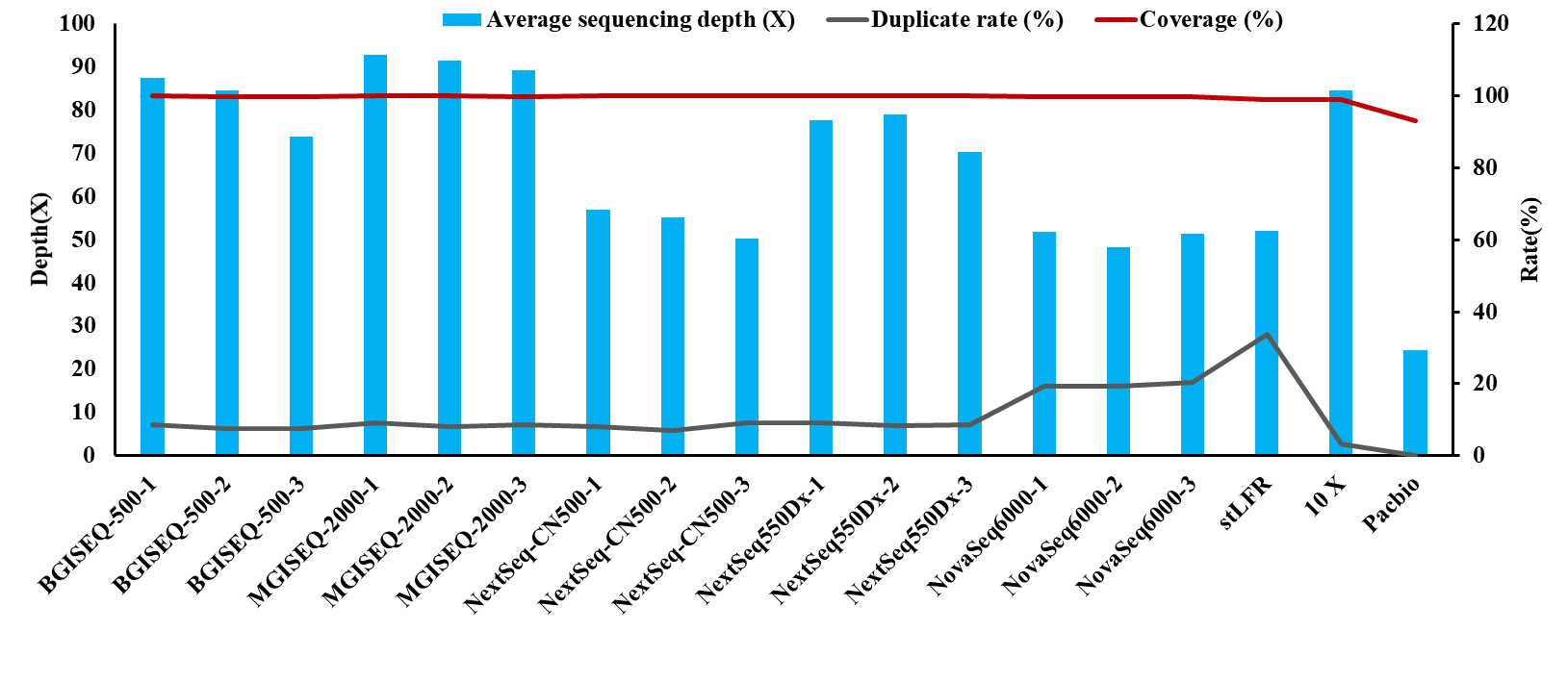

**Supplementary Figure S1. Sequencing depth, duplicate and mapping coverage.** For each platforms and duplicate sample, histograms indicate the sequencing depth, duplicate rate and coverage rate are represented by blue and gray line respectively.

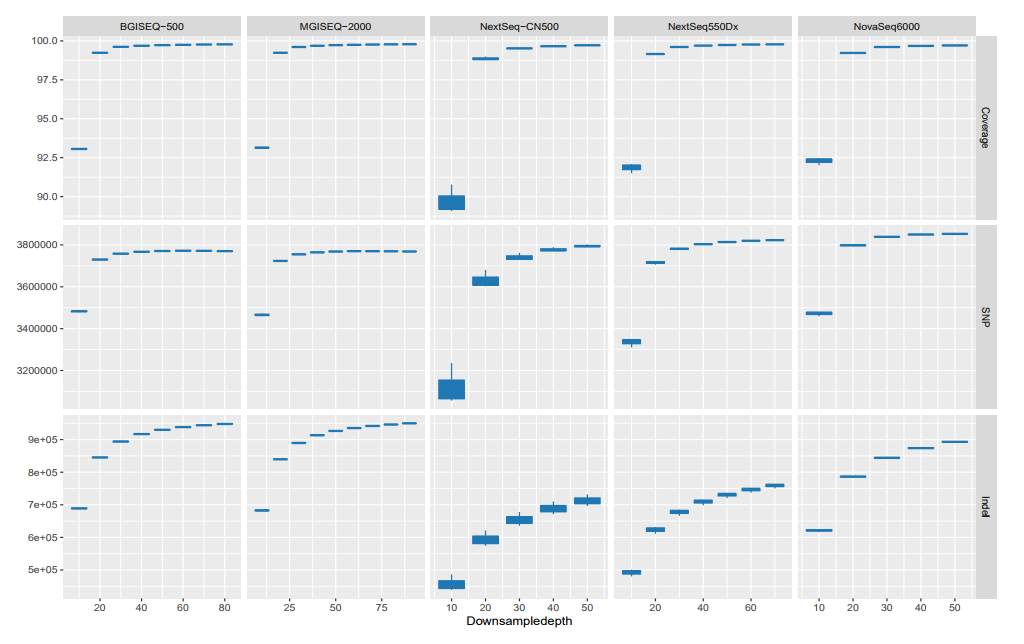

### Supplementary Figure S2. Saturate analysis for SNP, InDel and coverage for NGS platforms. Each windows shows average number for coverage, SNP and InDel

Numbers correspond to Sequencing depth from BGISEQ-500, MGISEQ-2000, NextSeq-CN500, NextSeq 550DX and NovaSeq6000 platforms.

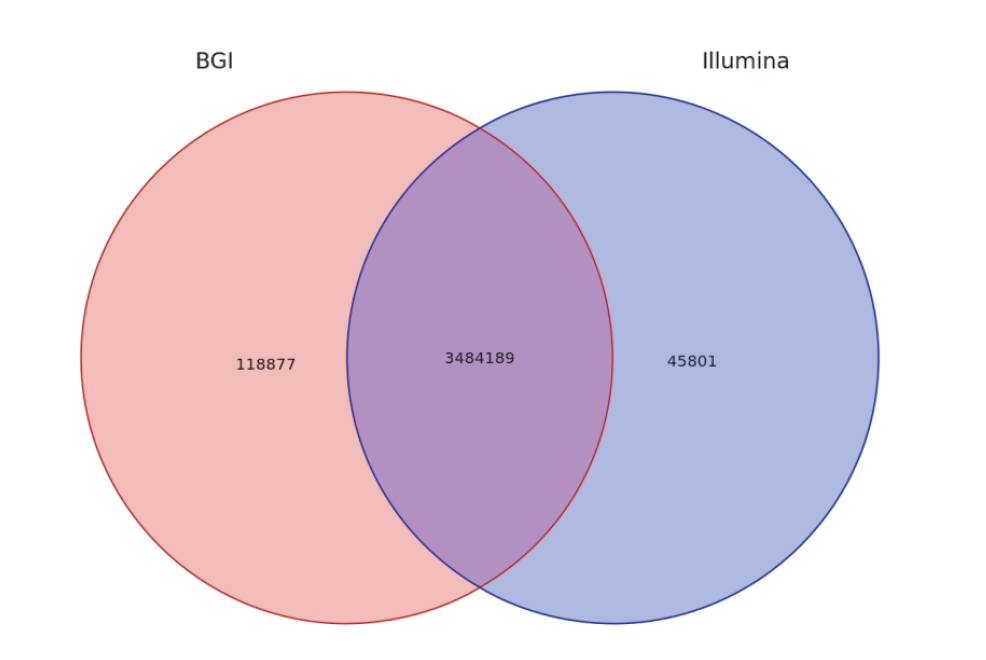

**Supplementary Figure S3**. **Consistency of SNPs from BGI and Illumina short sequence reads.** SNP consistency, supported by all samples for BGI and Illumina platforms are shown.

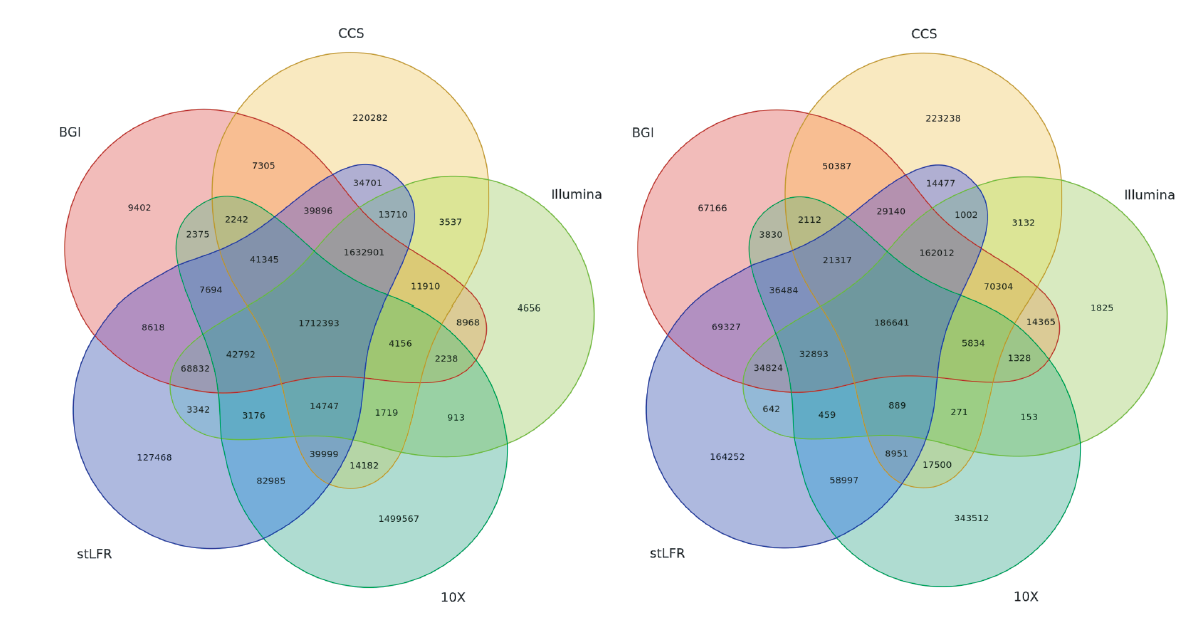

**A**

**B**

**Supplementary Figure S4. Consistence of SNPs and indels in UMR.** Consistency analysis: BGI regular NGS platforms, Illumina regular NGS platforms, two linked-read libraries, and PacBio CCS mode SNP(A) and indel(B) consistency analysis.

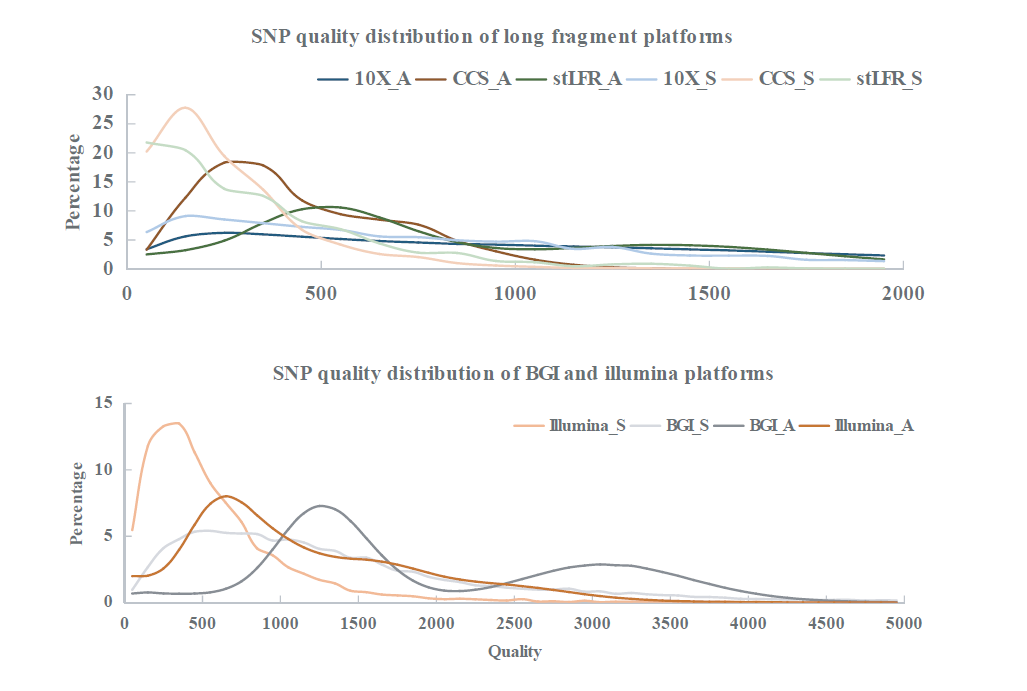

**A**

**B**

**Supplementary Figure S5. All and platform unique SNPs quality distribution for long fragment(A) and sort fragment platforms(B).** lines with tag “_A” indicates all SNP, “_S” means those unique.

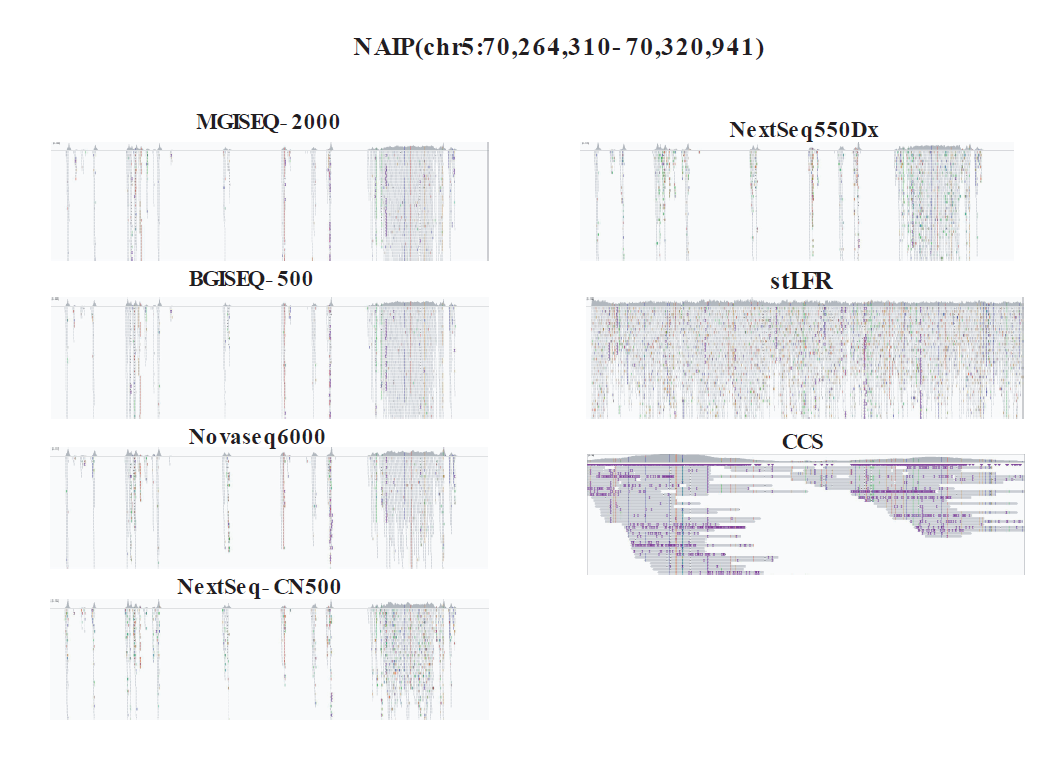

### Supplementary Figure S6. IGV views NAIP gene for each platforms.

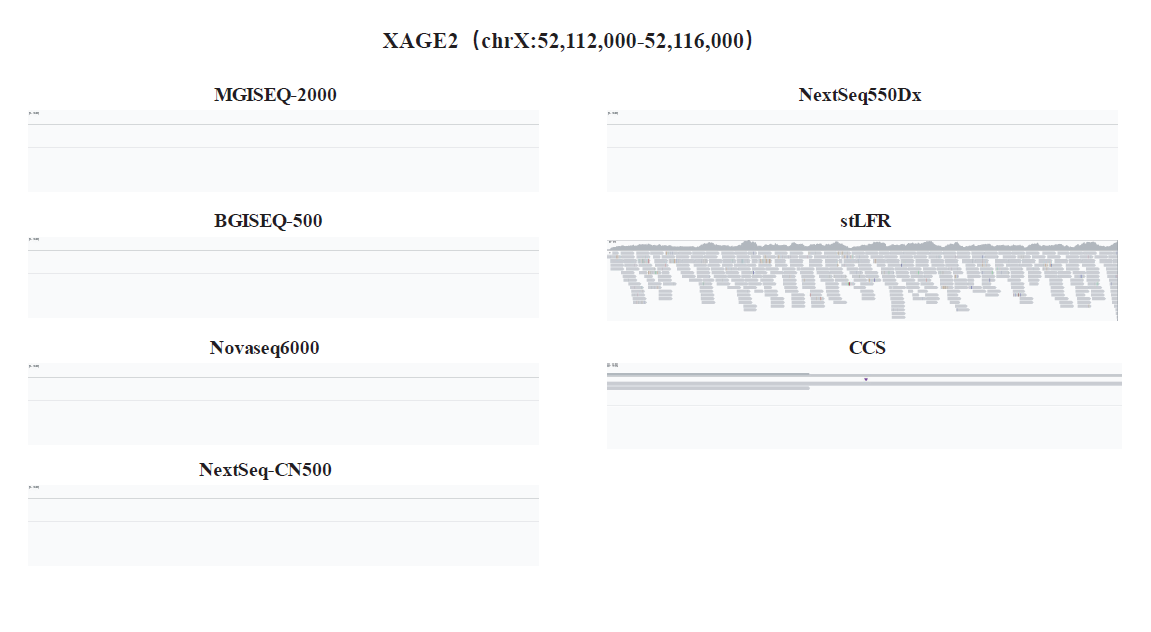

### Supplementary Figure S7. IGV views XAGE2 gene for each platforms.

**Supplementary Table S1-1. Data information of each platforms.**

|  | **BGISEQ-500** | | | **MGISEQ-2000** | | |
| --- | --- | --- | --- | --- | --- | --- |
|  | **1** | **2** | **3** | **1** | **2** | **3** |
| **Clean reads** | **2,835,787,762.00** | **2,705,531,402.00** | **2,366,532,060.00** | **3,025,576,552.00** | **2,947,464,316.00** | **2,895,152,490.00** |
| **Clean bases (Mb)** | **283,578.78** | **270,553.14** | **236,653.21** | **302,557.66** | **294,746.43** | **289,515.25** |
| **Mapping rate (%)** | **99.97** | **99.95** | **99.96** | **99.98** | **99.98** | **99.98** |
| **Unique rate (%)** | **87.89** | **88.99** | **88.91** | **87.28** | **88.44** | **87.84** |
| **Duplicate rate (%)** | **8.47** | **7.37** | **7.39** | **9.16** | **8.02** | **8.57** |
| **Mismatch rate (%)** | **0.22** | **0.21** | **0.19** | **0.25** | **0.28** | **0.28** |
| **Average sequencing depth (X)** | **87.50** | **84.52** | **73.86** | **92.77** | **91.57** | **89.25** |
| **Coverage (%)** | **99.93** | **99.92** | **99.92** | **99.93** | **99.93** | **99.92** |
| **Coverage at least 4X (%)** | **99.88** | **99.88** | **99.88** | **99.89** | **99.89** | **99.88** |
| **Coverage at least 10X (%)** | **99.82** | **99.81** | **99.79** | **99.82** | **99.82** | **99.81** |
| **Coverage at least 20X (%)** | **99.67** | **99.65** | **99.59** | **99.69** | **99.69** | **99.65** |
| **snp_number** | **3,731,347.00** | **3,722,674.00** | **3,712,120.00** | **3,685,587.00** | **3,696,424.00** | **3,707,393.00** |
| **Indel_number** | **958,346.00** | **961,643.00** | **954,785.00** | **958,497.00** | **962,494.00** | **958,369.00** |

**Supplementary Table S1-2. Data information of each platforms.**

|  | **NextSeq-CN500** | | | **NextSeq550Dx** | | |
| --- | --- | --- | --- | --- | --- | --- |
|  | **1** | **2** | **3** | **1** | **2** | **3** |
| **Clean reads** | **1,312,748,450.00** | **1,257,851,333.00** | **1,151,358,700.00** | **1,736,721,953.00** | **1,751,990,056.00** | **1,561,010,158.00** |
| **Clean bases (Mb)** | **196,912.27** | **188,677.70** | **172,703.80** | **262,245.01** | **259,721.10** | **231,618.38** |
| **Mapping rate (%)** | **100.00** | **99.68** | **99.68** | **99.77** | **99.78** | **99.81** |
| **Unique rate (%)** | **86.94** | **87.89** | **86.32** | **87.57** | **88.45** | **88.19** |
| **Duplicate rate (%)** | **7.92** | **7.02** | **9.08** | **9.05** | **8.29** | **8.57** |
| **Mismatch rate (%)** | **0.55** | **0.55** | **0.52** | **0.43** | **0.43** | **0.41** |
| **Average sequencing depth (X)** | **56.97** | **55.10** | **50.22** | **77.61** | **79.02** | **70.30** |
| **Coverage (%)** | **99.93** | **99.93** | **99.93** | **99.93** | **99.93** | **99.93** |
| **Coverage at least 4X (%)** | **99.86** | **99.86** | **99.85** | **99.89** | **99.89** | **99.88** |
| **Coverage at least 10X (%)** | **99.67** | **99.67** | **99.64** | **99.81** | **99.81** | **99.79** |
| **Coverage at least 20X (%)** | **98.67** | **98.49** | **98.16** | **99.59** | **99.60** | **99.53** |
| **snp_number** | **3,728,556.00** | **3,731,429.00** | **3,737,799.00** | **3,753,140.00** | **3,733,708.00** | **3,755,283.00** |
| **Indel_number** | **728,748.00** | **711,141.00** | **739,453.00** | **781,824.00** | **794,458.00** | **784,894.00** |

**Supplementary Table S1-3. Data information of each platforms.**

|  | **NovaSeq6000** | | | **Linked** | | **Pacbio** |
| --- | --- | --- | --- | --- | --- | --- |
|  | **1** | **2** | **3** | **stLFR** | **10X** | **CCS** |
| **Clean reads** | **1,289,631,254.00** | **1,199,680,727.00** | **1,294,029,261.00** | **2,507,844,034.00** | **2,004,349,222.00** | **6,409,943.00** |
| **Clean bases (Mb)** | **193,444.69** | **179,952.11** | **194,104.39** | **250,784.40** | **277,602.37** | **77,229.46** |
| **Mapping rate (%)** | **99.88** | **99.91** | **99.92** | **99.35** | **99.71** | **100.00** |
| **Unique rate (%)** | **77.47** | **77.60** | **76.49** | **65.05** | **95.38** | **99.82** |
| **Duplicate rate (%)** | **19.28** | **19.16** | **20.31** | **33.66** | **3.10** | **0.00** |
| **Mismatch rate (%)** | **0.24** | **0.26** | **0.24** | **0.00** | **0.00** | **0.00** |
| **Average sequencing depth (X)** | **51.83** | **48.24** | **51.32** | **51.97** | **84.70** | **24.40** |
| **Coverage (%)** | **99.90** | **99.91** | **99.91** | **98.86** | **98.90** | **93.18** |
| **Coverage at least 4X (%)** | **99.84** | **99.84** | **99.84** | **98.72** | **98.81** | **92.83** |
| **Coverage at least 10X (%)** | **99.71** | **99.71** | **99.73** | **98.34** | **98.63** | **90.18** |
| **Coverage at least 20X (%)** | **98.69** | **98.23** | **98.68** | **96.61** | **98.16** | **62.55** |
| **snp_number** | **3,751,598.00** | **3,716,984.00** | **3,734,579.00** | **3,874,599.00** | **3,472,522.00** | **3,795,024.00** |
| **Indel_number** | **944,658.00** | **943,428.00** | **944,173.00** | **822,307.00** | **721,170.00** | **797,206.00** |

| **Sample** | **of uncover_fragment** | **Length of uncover fragment** | **Ratio of uncover fragment** | **Max length** | **Mean length** | **1-100bp Rate** | **>100bp Rate** | **Repeat rate** | **LowComplexity**  **Rate** | **Mappability**  **Rate** |
| --- | --- | --- | --- | --- | --- | --- | --- | --- | --- | --- |
| **Common** | **51612** | **44414692** | **1.531361492** | **171822** | **860** | **43.88320546** | **56.11679454** | **28.06** | **3.55** | **63.58** |
| **BGISEQ-500-1** | **86531** | **56313296** | **1.941610064** | **191591** | **650** | **46.87915314** | **53.12084686** | **29.57** | **4.44** | **59.91** |
| **BGISEQ-500-2** | **88532** | **55809196** | **1.924229344** | **187034** | **630** | **48.99471378** | **51.00528622** | **29.61** | **4.55** | **59.69** |
| **BGISEQ-500-3** | **91721** | **56667668** | **1.953828355** | **187034** | **617** | **49.5698913** | **50.4301087** | **29.7** | **4.57** | **59.36** |
| **MGISEQ-2000-1** | **84899** | **55945722** | **1.928936585** | **187034** | **658** | **46.66839421** | **53.33160579** | **29.52** | **4.38** | **60.12** |
| **MGISEQ-2000-2** | **86376** | **55242444** | **1.904688464** | **186504** | **639** | **48.74270631** | **51.25729369** | **29.47** | **4.45** | **60.07** |
| **MGISEQ-2000-3** | **87463** | **55712134** | **1.920882771** | **186480** | **636** | **48.52223226** | **51.47776774** | **29.74** | **4.58** | **59.72** |
| **NextSeq550Dx-1** | **83478** | **50647655** | **1.746266045** | **191340** | **606** | **57.32887707** | **42.67112293** | **28.18** | **4.78** | **60.51** |
| **NextSeq550Dx-2** | **82407** | **49566962** | **1.709005139** | **186428** | **601** | **58.15646729** | **41.84353271** | **28.06** | **4.85** | **60.39** |
| **NextSeq550Dx-3** | **89348** | **49892509** | **1.720229581** | **191283** | **558** | **61.03662085** | **38.96337915** | **28.07** | **4.93** | **60.01** |
| **NextSeq-CN500-1** | **114603** | **58055997** | **2.001696155** | **191434** | **506** | **59.23754177** | **40.76245823** | **28.96** | **5.58** | **58.48** |
| **NextSeq-CN500-2** | **113902** | **58131009** | **2.004282472** | **191558** | **510** | **59.32029288** | **40.67970712** | **28.91** | **5.44** | **58.45** |
| **NextSeq-CN500-3** | **103920** | **55262351** | **1.905374832** | **191311** | **531** | **60.31947652** | **39.68052348** | **28.62** | **5.63** | **58.71** |
| **NovaSeq6000-1** | **66490** | **54766189** | **1.888267804** | **186495** | **823** | **38.23432095** | **61.76567905** | **29.29** | **4.31** | **59.78** |
| **NovaSeq6000-2** | **65357** | **53158741** | **1.832845063** | **187034** | **813** | **41.36358768** | **58.63641232** | **29.05** | **4.22** | **59.73** |
| **NovaSeq6000-3** | **64218** | **53005900** | **1.827575301** | **186471** | **825** | **40.68018313** | **59.31981687** | **29.12** | **4.32** | **59.77** |

**Supplementary Table S2. Statistics of NGS uncovered regions.**

**Supplementary Table S3. Phasing statistic for PacBio SMRT CCS, stLFR and 10X Genomics Chromium.**

|  | **CCS** | **stLFR** | **10X** |
| --- | --- | --- | --- |
| **Length(bp)** | **30403236** | **32519955** | **29020167** |
| **Percentage(%)** | **68.53** | **73.3** | **65.41** |
| **UTR3** | **61** | **155** | **193** |
| **UTR5** | **13** | **44** | **59** |
| **Downstream** | **348** | **421** | **442** |
| **exonic** | **397** | **441** | **511** |
| **Exonic;Splicing** | **1** | **1** | **1** |
| **Intergenic** | **16480** | **29512** | **31715** |
| **NcRNA_intronic** | **5220** | **6489** | **7112** |
| **NcRNA_exonic** | **416** | **517** | **557** |
| **NcRNA_intronic** | **3100** | **5866** | **5784** |
| **NcRNA_splicing** | **2** | **8** | **7** |
| **NcRNA_splicing** | **1** | **3** | **3** |
| **Upstream** | **471** | **448** | **490** |
| **Upstream;Downstream** | **14** | **7** | **20** |
